# Genome-wide association study identifies genetic factors that modify age at onset in Machado-Joseph disease

**DOI:** 10.1101/834754

**Authors:** Fulya Akçimen, Sandra Martins, Calwing Liao, Cynthia V. Bourassa, Hélène Catoire, Garth A. Nicholson, Olaf Riess, Mafalda Raposo, Marcondes C. França, João Vasconcelos, Manuela Lima, Iscia Lopes-Cendes, Maria Luiza Saraiva-Pereira, Laura B. Jardim, Jorge Sequeiros, Patrick A. Dion, Guy A. Rouleau

**Affiliations:** Department of Human Genetics, McGill University, Montréal, QC, Canada; Montreal Neurological Institute and Hospital, McGill University, Montréal, QC, Canada; i3S – Instituto de Investigação e Inovação em Saúde, Universidade do Porto, Portugal; IPATIMUP – Institute of Molecular Pathology and Immunology of the University of Porto, Portugal; Department of Neurology and Neurosurgery, McGill University, Montréal, QC, Canada; University of Sydney, Department of Medicine, Concord Hospital, Australia; Institute of Medical Genetics and Applied Genomics, University of Tuebingen, Tuebingen, Germany; Faculdade de Ciências e Tecnologia, Universidade dos Açores e Instituto de Biologia Molecular e Celular (IBMC), Instituto de Investigação e Inovação em Saúde (i3S), Universidade do Porto, Portugal; Department of Neurology, Faculty of Medical Sciences, UNICAMP, Campinas, SP, Brazil; School of Medical Sciences, Department of Medical Genetics and Genomic Medicine, University of Campinas (UNICAMP), Campinas, SP, Brazil; The Brazilian Institute of Neuroscience and Neurotechnology (BRAINN), Campinas, SP, Brazil; Departamento de Neurologia, Hospital do Divino Espírito Santo, Ponta Delgada, Portugal; Medical Genetics Service, Hospital de Clínicas de Porto Alegre (HCPA), Porto Alegre, Brazil; Depto. de Bioquímica – ICBS, Universidade Federal do Rio Grande do Sul (UFRGS); Depto de Medicina Interna, Universidade Federal do Rio Grande do Sul (UFRGS), Porto Alegre, Brazil; Institute for Molecular and Cell Biology (IBMC), Universidade do Porto, Porto, Portugal; Instituto de Ciências Biomédicas Abel Salazar (ICBAS), Universidade do Porto, Portugal

**Keywords:** Machado-Joseph disease, *ATXN3*, MJD/SCA3, age at onset, modifier, GWAS

## Abstract

Machado-Joseph disease (MJD/SCA3) is the most common form of dominantly inherited ataxia worldwide. The disorder is caused by an expanded CAG repeat in the *ATXN3* gene. Past studies have revealed that the length of the expansion partly explains the disease age at onset (AO) variability of MJD, which is confirmed in this study. Using a total of 786 MJD patients from five different geographical origins, a genome-wide association study (GWAS) was conducted to identify additional AO modifying factors that could explain some of the residual AO variability. We identified nine suggestively associated loci (*P* < 1 × 10^−5^). These loci were enriched for genes involved in vesicle transport, olfactory signaling, and synaptic pathways. Furthermore, associations between AO and the *TRIM29* and *RAG* genes suggests that DNA repair mechanisms might be implicated in MJD pathogenesis. Our study demonstrates the existence of several additional genetic factors, along with CAG expansion, that may lead to a better understanding of the genotype-phenotype correlation in MJD.

## Introduction

Machado-Joseph disease, also known as spinocerebellar ataxia type 3 (MJD/SCA3), is an autosomal dominant neurodegenerative disorder that is characterized by progressive cerebellar ataxia and pyramidal signs, which can be associated with a complex clinical picture and includes extrapyramidal signs or amyotrophy [1, 2]. MJD is caused by an abnormal CAG trinucleotide repeat expansion in exon 10 of the ataxin-3 gene (*ATXN3*), located at 14q32.1. Deleterious expansions consensually contain 61 to 87 CAG repeats, whereas wild type alleles range from 12 to 44 [2].

As with other diseases caused by repeat expansions, such as Huntington’s disease (HD) and other spinocerebellar ataxias, there is an inverse correlation between expanded repeat size and the age at which pathogenesis leads to disease onset [3]. Depending on the cohort structure, the size of the repeat expansion explains 55 to 70% of the age at onset (AO) variability in MJD, suggesting the existence of additional modifying factors [3,4]. Although several genetic factors have been proposed as modifiers, such as CAG repeat size of normal *ATXN3* (SCA3), *HTT* (HD), *ATXN2* (SCA2) and *ATN1* (DRPLA) alleles, *APOE* status, and expression level of *HSP40* [4,5,6], these were not replicated by subsequent studies [7, 8]. Since CAG tract profile and allelic frequencies of the potential modifier loci can have unique characteristics in different populations, large collaborative studies are required to identify genetic modifiers in MJD, as well as replicate the findings of such studies [8].

Previously, Genetic Modifiers of Huntington’s Disease (GeM-HD) Consortium carried out a GWA approach of HD individuals to reveal genetic modifiers of AO in HD [9,10]. A total of eleven [9] and fourteen loci [10] were found to be associated with residual age at HD onset. In the present study, we performed the first GWAS to identify some possible genetic modifiers of AO in MJD. First, we assessed the relationship between AO and size of the expanded (CAG_exp_) and normal (CAG_nor_) alleles, biological sex and geographical origin. Next, we determined a residual AO for each subject, which is the difference between the measured AO and the predicted/estimated AO from expanded CAG repeat size alone. Using the residuals as a quantitative phenotype for a GWAS, we looked for genetic factors that modulate AO in MJD.

## Methods

### Study subjects

A total of 786 MJD patients from five distinct geographical origins (Portugal, Brazil, North America, Germany and Australia) were included in the present study. The overall average age at onset (standard deviation) was 38 (± 1.82) years, with a 1:1 male to female ratio. All subjects provided informed consent, and the study was approved by the respective institutional review boards. Detailed cohort demographics are shown in Supplementary Table 1.

### Assessment of the *ATXN3* CAG repeat length

A singleplex polymerase chain reaction was performed to determine the length of the CAG_exp_ and CAG_nor_ alleles at exon 10 of *ATXN3* [11]. The final volume for each assay was 10 μL: 7.5 ng of gDNA, 0.2 μM of each primer, 5 μL of Taq PCR Master Mix Kit Qiagen®, 1 μL of Q-Solution from Qiagen® and H_2_O. Fragment length analysis was done using ABIPrism 3730×l sequencer (Applied Biosystems®, McGill University and Genome Québec Innovation Centre) and GeneMapper software [12]. A stepwise regression model was performed to assess the correlation between AO and CAG_exp_ size, as well as gender, origin, CAG_nor_ size, and interaction between these variables. Residual AO was calculated for each subject by subtracting individual’s expected AO based upon CAG_exp_ size from actual AO, to be used as the primary phenotype for following genetic approach.

### Genotyping, quality control and imputation

Samples were genotyped using the Global Screening Array v.1.0 from Illumina (636,139 markers). Sample-based (missingness, relatedness, sex, and multidimensional scaling analysis) and SNP-based quality assessments (missingness, Hardy-Weinberg equilibrium, and minor allele frequency) were conducted using PLINK version 1.9 [13]. In sample level QC, samples were excluded with one or more of the following: high missingness (missingness rate > 0.05), close relationship (pi-hat value > 0.2), discrepancy between genetically-inferred sex and reported sex, population outliers (deviation ≥ 4 SD from the population mean in multidimensional scaling analysis). All SNPs were checked for marker genotyping call rate (> 98%), minor allele frequency (MAF) > 0.05, and HWE (p-value threshold = 1.0 × 10^−5^).

Phasing and imputation were performed using SHAPEIT [14] and PBWT [15] pipelines, implemented on the Sanger Imputation Service [16]. Haplotype Reference Consortium (HRC) reference panel r1.1 containing 64,940 human haplotypes at 40,405,505 genetic markers were used as the reference panel. Imputed variants with an allele count of 30 (MAF > 0.02), an imputation quality score above 0.3 and an HWE p-value of > 1.0 × 10^−5^ were included for subsequent analysis.

### Genome-wide association analysis

A genome-wide linear mixed model based association analysis was conducted using GCTA version 1.91.7 [17]. Residual AO was modelled as a function of minor allele count of the test SNP, sex, and the first three principal components based on the scree plot (Supplementary Figure 1). The --mlma-loco option, which takes into account the difference in allele frequency between populations, was used to control for population structure. QQ plots and Manhattan plots were generated in FUMA v.1.3.4 [18]. Regional association plots were generated using LocusZoom [19] (Supplementary Figure 3).

### Functional annotation of SNPs

Genomic risk loci were defined using SNP2GENE function implemented in FUMA. Independent suggestive SNPs (*P* < 1 × 10^−5^) with a threshold of r^2^ < 0.6 were selected within a 250 kb window. The UK Biobank release 2 European population consisting of randomly selected 10,000 subjects was used as the reference population panel. The ANNOVAR [20] categories and combined annotation-dependent depletion (CADD) [21] scores were obtained from FUMA for functional annotation. Functionally annotated variants were mapped to genes based on genomic position using FUMA positional mapping tool.

### Pathway analysis

To identify known biological pathways and gene sets at the associated loci, an enrichment approach was applied using public datasets containing Gene Ontology (GO, http://geneontology.org), the Kyoto Encyclopaedia of Genes and Genomes (KEGG, https://www.genome.jp/kegg) and Reactome (https://reactome.org) pathways. The primary enrichment analysis was performed using the i-GSEA4GWAS v2 [22]. It uses a candidate list of a genome-wide set of genes mapped within the SNP loci and ranks them based on the strength of their association with the phenotype. Genes were mapped within 20 kb up or downstream of the SNPs with a *P*<0.05. Gene and pathway sets meeting a false discovery rate *(FDR)*-corrected *q*-value < 0.05 were regarded as significantly associated with high confidence, and *q*-value < 0.25 was regarded to be possibly associated with the phenotype of interest. We performed a secondary gene-based association test using the Versatile Gene-based Association Study (VEGAS) [23] algorithm that controls the number of SNPs in each gene and the linkage disequilibrium (LD) between these SNPs using the HapMap European population. As a third algorithm to identify enriched pathways, we used Pathway Scoring Algorithm (PASCAL) [24], which controls for potential bias from gene size, SNP density, as well as LD. ClueGO [25] and CluePedia [26] plug-ins in Cytoscape were employed to visualize identified pathways and their clustering.

## Results

### The inverse correlation between CAG_exp_ and age at onset

In the first phase of the study, the expanded *ATXN3*-CAG repeat lengths of 786 MJD patients were assessed. The mean (SD) CAG_exp_ size were Australia: 68.2 (±3.3), Brazil: 74.3 (3.9), Germany: 72.9 (±3.6), North America: 73 (±4.3) and Portugal: 72 (±4.0). Next, the relationship between AO and CAG_exp_ size, CAG_nor_ size, sex and ethnicity was examined (Supplementary Table 1). The previously observed negative correlation between *ATXN3* CAG_exp_ size and AO [3] was confirmed (Pearson’s correlation coefficient R^2^ = 0.62) (Figure 1). The CAG_nor_ size (P = 0.39), sex (P = 0.02) and geographic origin (P [Brazil] = 0.38, P [Germany] = 0.38, P [North America] = 0.33, P [Portugal] = 0.29) were not significant and their addition had little contribution to the model (ΔR^2^ = 0.0072). Residual AO for each sample was calculated and used as a quantitative phenotype to identify the modifiers of AO. The distribution of residual AO was close a theoretical normal distribution (Figure 1).

**Figure 1.**
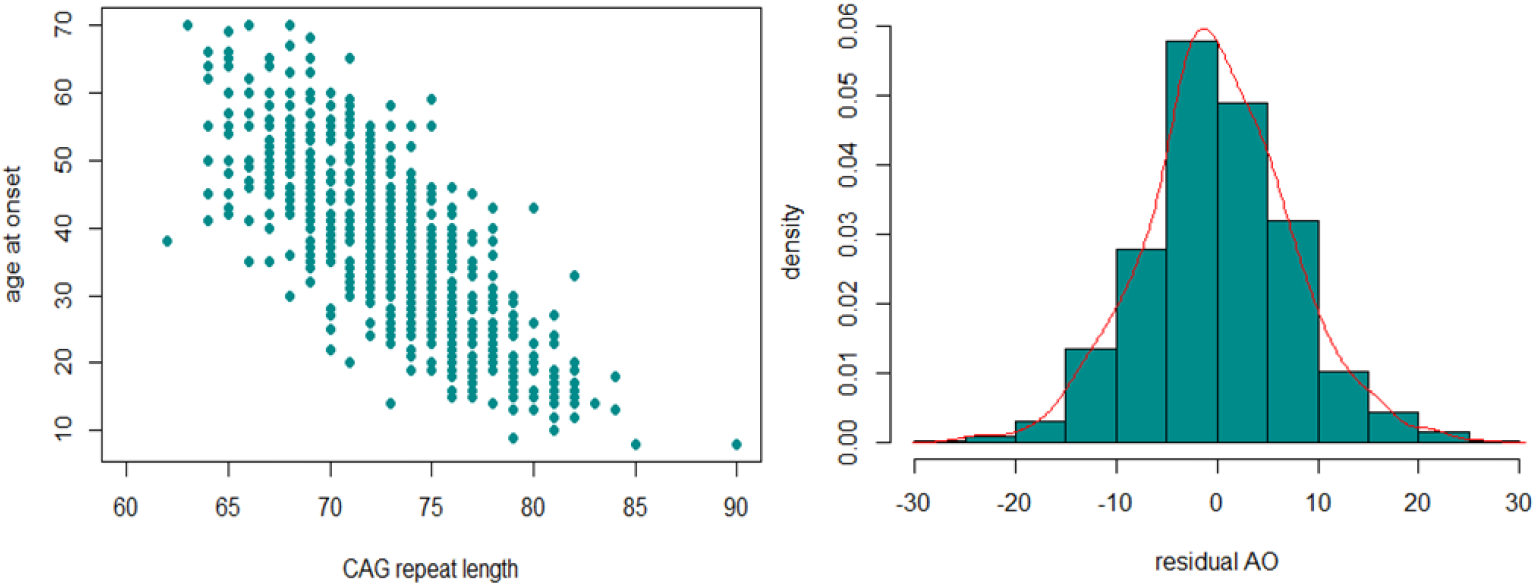
The inverse correlation between CAG_exp_ and AO (left) and the distribution of residual AO (right) observed in our MJD cohort.

### Genome-wide association study

After post-imputation quality assessments, a total of 700 individuals with genotyping information for 6,716,580 variants remained for GWAS. The resulting Manhattan plots and quantile-quantile (QQ) plots are shown in Figure 2. The genomic inflation factor was close to one (λ = 0.98), indicating the p-values were not inflated. No association signal was identified meeting genome-wide significance (P < 5 × 10^−8^, the genome-wide Bonferroni-corrected significance threshold); however, genome-wide suggestive associations (P < 1 × 10^−5^) with 204 variants across 9 loci were identified (Supplementary Table 3). The most significantly associated SNP at each locus are shown in Table 1. Positional gene mapping aligned SNPs to 17 genes by their genomic location. Fourteen of the 204 variants had a Combined Annotation Dependent Depletion (CADD)-PHRED score higher than the suggested threshold for deleterious SNPs (12.37), arguing the given loci have a functional role [27].

**Table 1.**
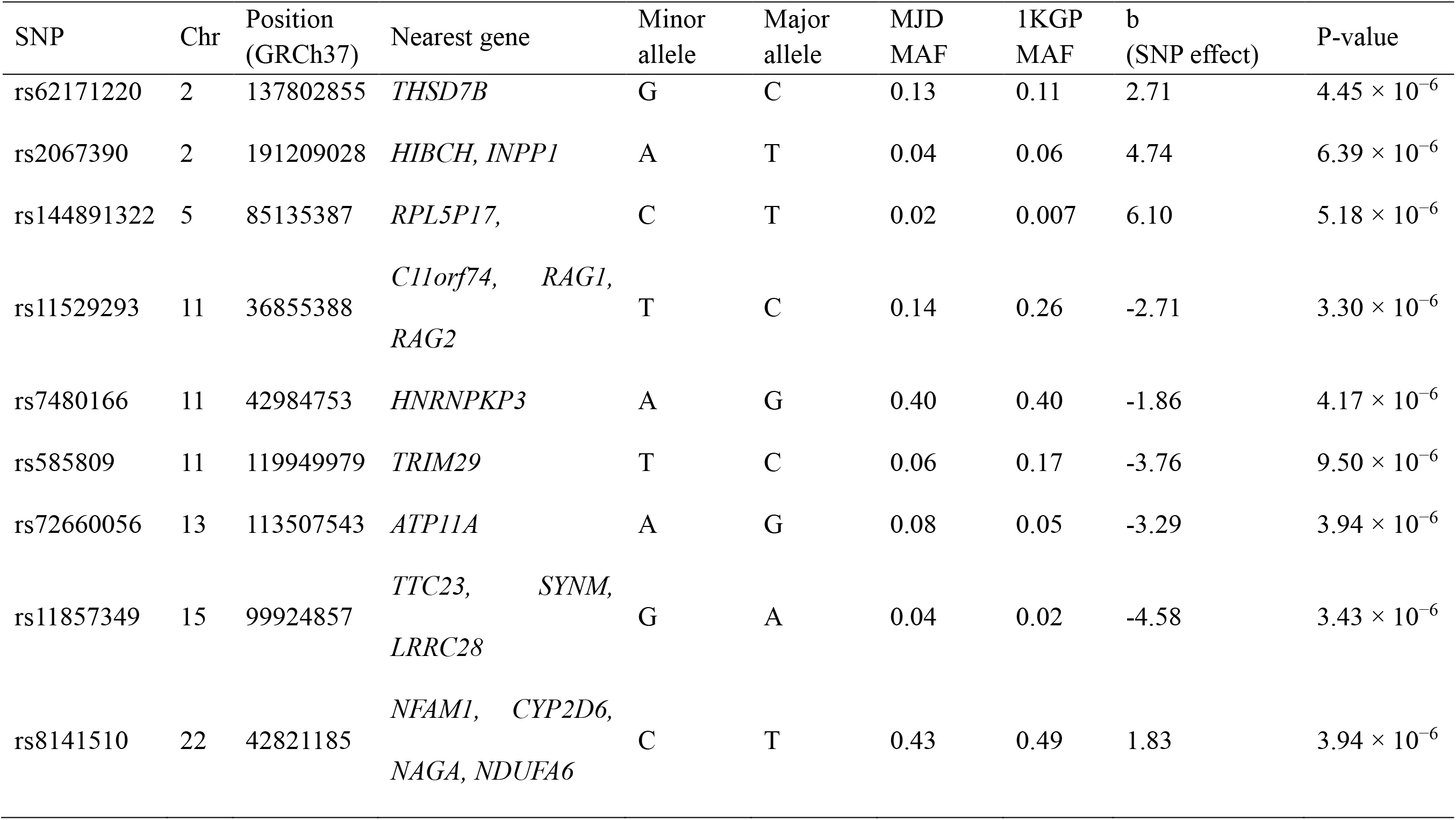
Suggestive loci associated with residual age at onset in MJD. Chr: chromosome, MAF: minor allele frequency, 1KGP: 1000 Genomes Project

**Figure 2.**
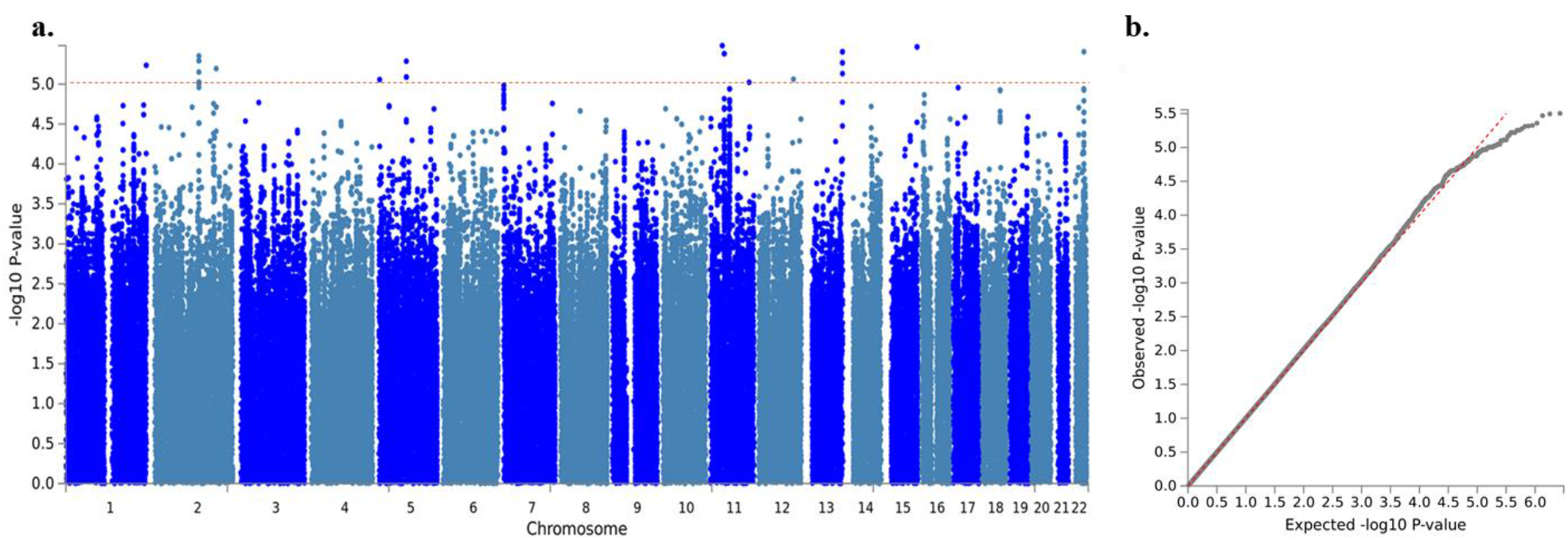
Manhattan plot (a) and QQ plot (b) of the GWAS for residual AO of MJD. Imputed using the HRC panel, 6,716,580 variants that passed QC are included in the plot. The x-axis shows the physical position along the genome. The y-axis shows the −log_10_(p-value) for association. The red line indicates the level of genome-wide suggestive association (*P* = 1 × 10^−5^).

### Interaction analysis between CAG_exp_ and SNP genotype

To assess a possible interaction between CAG_exp_ size and the variants identified, each of the nine variants was added to the initial linear regression, modelling AO as a function of CAG_exp_ size, SNP, sex, the first three principal components, CAG_nor_ size, and interaction of SNP:CAG_exp_. Association of each independent SNP with AO revealed nominally significant p-values. Among the nine variants, only rs585809 (mapped to *TRIM29*) had a significant interaction with CAG_exp_ (P = 0.01), suggesting that rs585809 might modulate AO through this epistatic interaction on CAG_exp_.

### Association of HD-AO modifier variants in MJD

Association of previously identified HD-AO modifier loci in MJD were assessed. Among the 25 HD-AO modifier variants in 17 loci, a total of 18 variants (MAF > 0.02) in 12 loci were tested in this study (Supplementary Table 4). None of these HD-AO modifiers reached the genome-wide suggestive threshold. However, two variants rs144287831 (*P* = 0.02, effect size = − 0.98) and rs1799977 (*P* = 0.02, effect size = − 0.98) in the *MLH1* locus were found to be nominally associated with a later AO in MJD.

### Pathway and gene-set enrichment analysis

A gene-set enrichment and pathway analysis was conducted using i-GSEA4GWAS. Various approaches and algorithms are currently in use to conduct similar analyses. To be able to make better comparisons with other studies that may use different approaches, we performed a secondary gene-set enrichment and pathway analysis using the VEGAS2 and PASCAL software (Supplementary Tables 5-7). We also used these results for replication purposes in our own study. A total of 13 overrepresented pathways were found, after FDR-multiple testing correction (q-value < 0.05) in the primary GSEA analysis and replicated using at least one of the secondary gene-set enrichment algorithms (Table 2). Overall, the most significantly enriched gene-sets and pathways were vesicle transport, olfactory signaling, and synaptic pathways. Visualization and clustering of pathways are shown in Figure 3.

**Figure 3.**
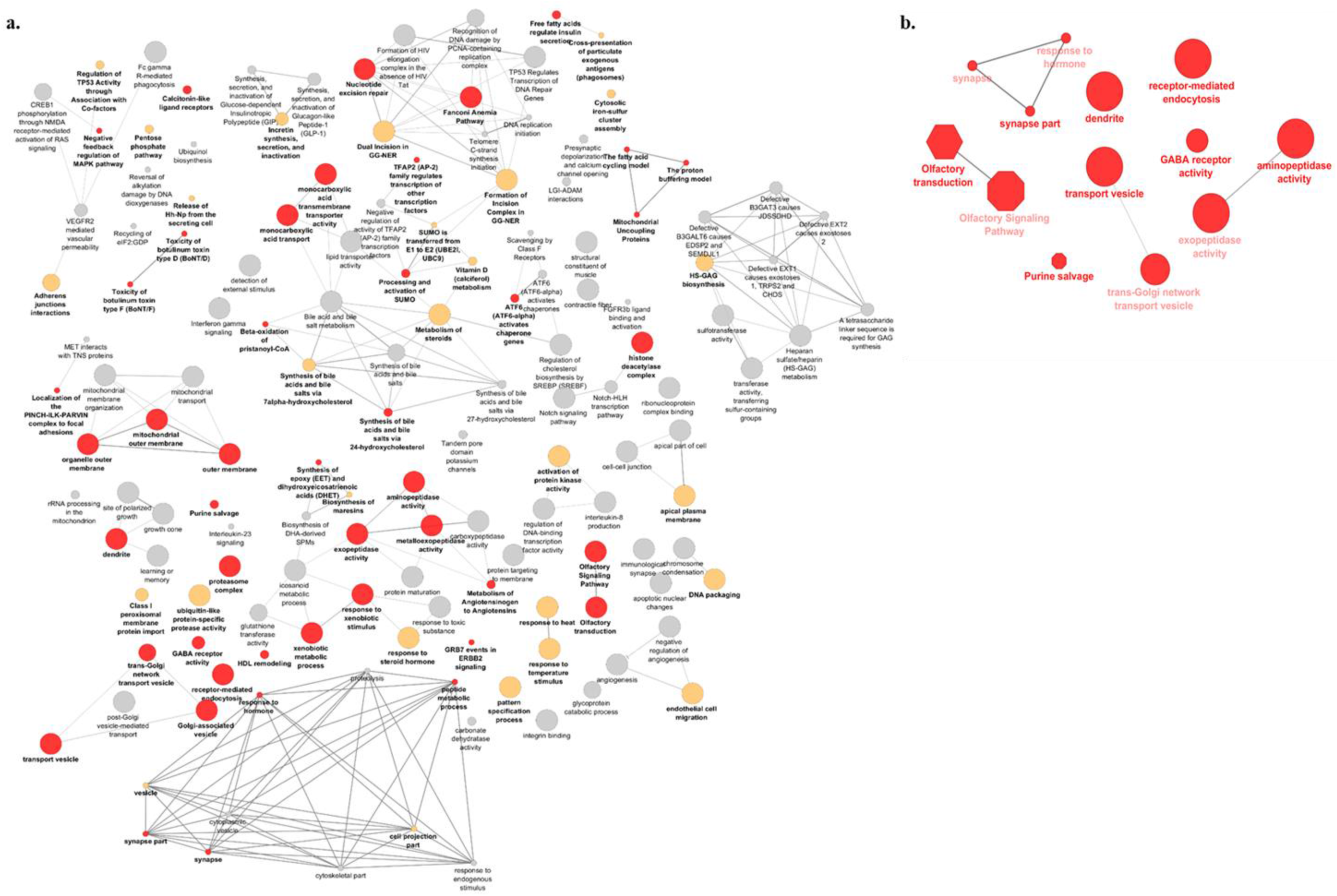
Visualization of the gene-sets and pathways enriched in primary GSEA analysis (a) and replicated in VEGAS and PASCAL (b). The size of the nodes corresponds to the number of the genes associated with a term. The significance is represented by the color of the nodes (*P* < 0.05, 0.05 < *P* < 0.1 and *P* > 0.1 are represented by red, yellow and gray, respectively).

**Table 2.** Pathways significant after multiple-correction (q < 5 × 10^−2^) in the primary GSEA analysis and replicated using at least one of the secondary gene-set enrichment algorithms. NA means that the pathway was not enriched by at least two significant genes in VEGAS.

| Pathway | Description | p-value<br>(GSEA) | q-value<br>(GSEA) | p-value<br>(VEGAS) | permuted<br>p-value (VEGAS) | p-value<br>(PASCAL) |
| --- | --- | --- | --- | --- | --- | --- |
| GO:0030133 | transport vesicle | $< 1.0 \times 10^{-3}$ | $8.20 \times 10^{-3}$ | $6.15 \times 10^{-40}$ | $4.46 \times 10^{-1}$ | $6.70 \times 10^{-3}$ |
| KEGG:04740 | olfactory transduction | $< 1.0 \times 10^{-3}$ | $8.30 \times 10^{-3}$ | NA | NA | $3.89 \times 10^{-4}$ |
| R-HSA:381753 | olfactory signaling pathway | $< 1.0 \times 10^{-3}$ | $8.80 \times 10^{-3}$ | $1.10 \times 10^{-27}$ | $7.71 \times 10^{-1}$ | $2.51 \times 10^{-4}$ |
| GO:0044456 | synapse part | $< 1.0 \times 10^{-3}$ | $9.30 \times 10^{-3}$ | $1.25 \times 10^{-182}$ | $< 1.0 \times 10^{-6}$ | $< 1.0 \times 10^{-7}$ |
| R-HSA:74217 | purine salvage | $< 1.0 \times 10^{-3}$ | $1.06 \times 10^{-2}$ | $1.06 \times 10^{-2}$ | $2.15 \times 10^{-1}$ | $6.48 \times 10^{-3}$ |
| GO:0045202 | synapse | $< 1.0 \times 10^{-3}$ | $1.15 \times 10^{-2}$ | $1.15 \times 10^{-2}$ | $< 1.0 \times 10^{-6}$ | $< 1.0 \times 10^{-7}$ |
| GO:0004177 | aminopeptidase activity | $< 1.0 \times 10^{-3}$ | $1.50 \times 10^{-2}$ | $1.50 \times 10^{-2}$ | $3.41 \times 10^{-1}$ | $1.24 \times 10^{-2}$ |
| GO:0008238 | exopeptidase activity | $< 1.0 \times 10^{-3}$ | $1.80 \times 10^{-2}$ | $1.80 \times 10^{-2}$ | $2.80 \times 10^{-2}$ | $8.31 \times 10^{-3}$ |
| GO:0006898 | receptor mediated endocytosis | $< 1.0 \times 10^{-3}$ | $2.25 \times 10^{-2}$ | $2.25 \times 10^{-2}$ | $2.03 \times 10^{-1}$ | $6.64 \times 10^{-3}$ |
| GO:0016917 | GABA receptor activity | $< 1.0 \times 10^{-3}$ | $2.26 \times 10^{-2}$ | $2.26 \times 10^{-2}$ | $1.30 \times 10^{-4}$ | $2.30 \times 10^{-5}$ |
| GO:0030140 | trans Golgi network transport vesicle | $< 1.0 \times 10^{-3}$ | $2.36 \times 10^{-2}$ | $2.36 \times 10^{-2}$ | $2.80 \times 10^{-2}$ | $1.28 \times 10^{-1}$ |
| GO:0009725 | response to hormone stimulus | $< 1.0 \times 10^{-3}$ | $2.73 \times 10^{-2}$ | $2.73 \times 10^{-2}$ | $1.32 \times 10^{-1}$ | $1.30 \times 10^{-4}$ |
| GO:0030425 | Dendrite | $< 1.0 \times 10^{-3}$ | $3.86 \times 10^{-2}$ | $3.86 \times 10^{-2}$ | $< 1.0 \times 10^{-6}$ | $< 1.0 \times 10^{-7}$ |

## Discussion

Using five cohorts from different geographical origins, we performed the first GWAS to examine the presence of genetic factors that could modify AO in MJD. We identified a total of nine loci that were potentially associated with either an earlier or later AO. Concomitantly, we confirmed the previously observed negative correlation between CAG_exp_ and AO [3]. It was shown previously that normal *ATXN3* allele (CAG_nor_) had a significant influence on AO of MJD [28]; however, several studies did not replicate this effect [6,8]. Indeed, we did not observe an association between CAG_nor_ and AO. However, it had little contribution to our model, with a minor difference in the correlation coefficient (ΔR^2^ = 0.0012).

In our GWAS, the strongest signal is for the rs11529293 variant (*P* = 3.30 × 10^−6^) within the *C11orf72* and *RAG* loci at 11p12. Within this locus, two *RAG* genes, recombination-activating genes *RAG1* and *RAG2*, were shown to be implicated in DNA damage response and DNA repair machineries [29,30]. The rs585809 variant, which was mapped to the *TRIM29* gene, was found to interact with CAG_exp_, suggesting that it might have an effect on AO through this interaction. Both *RAG* and *TRIM29* loci were identified as AO-hastening modifiers. *TRIM29* encodes for tripartite motif protein 29, which is implicated in mismatch repair and double strand breaks pathways [31,32]. TRIM29 is involved both upstream and downstream of these pathways, in the regulation of DNA repair proteins into chromatin by mediating the interaction between them. One of these DNA repair proteins is MLH1, which is implicated in mismatch repair complex [32]. Previously, the *MLH1* locus was identified as an AO modifier in another neurodegenerative disease caused by CAG repeat expansion, Huntington’s disease [9,10,33]. Additionally, in a genome-wide genetic screening study, MLH1-knock out was shown to modify the somatic expansion of the CAG repeat and slow the pathogenic process in HD mouse model [34]. Overall, the association of *TRIM29* and *RAG* loci suggests that DNA repair mechanisms may be implicated in the alteration of AO of MJD, as well as HD, and may have a role in the pathogenesis of other CAG repeat diseases. Interestingly, in a previous study, we found variants in three transcription-coupled repair genes (ERCC6, RPA, and CDK7) associated with different CAG instability patterns in MJD [35].

We identified gene-sets enriched in olfactory signaling, vesicle transport, and synaptic pathways. Olfactory dysfunction is one of the main non-motor symptoms that was already described in patients with MJD [36,37]. In a previous study, transplantation of olfactory ensheathing cells, which are specialized glial cells of the primary olfactory system, were found to improve motor function in an MJD mice model, and were suggested as a novel potential strategy for MJD treatment [38]. Vesicle transport and synaptic pathways were also implicated in MJD, as well as in other neurodegenerative diseases [39,40]. An interruption of synaptic transmission caused by an expanded polyglutamine repeat and mutant ataxin-3 aggregates were shown in *Drosophila* and *Caenorhabditis elegans* models of MJD. Therefore, the interaction between synaptic vesicles and mutant aggregates supports the role of synaptic vesicle transport in the pathogenesis of MJD [41,42]. Overall, we suggest that these gene-sets and pathways might construct a larger molecular network that could modulate the AO in MJD.

In summary, our study identified nine genetic loci that may modify the AO of MJD. Identification of *TRIM29* and *RAG* genetic variants, as well as our gene-set enrichment analyses, implicated DNA repair, olfactory signaling, synaptic, and vesicle transport pathways in the pathogenesis of MJD. Although we used different cohorts from five distinct geographical ethnicities, a replication study in similar or additional populations would add valuable evidence to support our findings.

## Description of Supplemental Data

Supplemental Data include three figures and seven tables

## Acknowledgements

The authors thank the participants for their contribution to the study. The authors would like to thank Jay P. Ross, Faezeh Sarayloo, Zoe Schmilovich and S. Can Akerman for their assistance in reviewing the manuscript and scientific content. FA was funded by the Fonds de Recherche du Québec–Santé. SM is funded by FCT (CEECIND/00684/2017) and by NORTE-01-0145-FEDER-000008, supported by Norte Portugal Regional Programme (NORTE 2020), under the PORTUGAL 2020 Partnership Agreement, through the European Regional Development Fund (ERDF). FM and LI are funded by Fundaçao de Amparo a Pesquisa do Estado de São Paulo (FAPESP, 2013/07559-3). MLSP and LBJ were funded by Conselho Nacional de Desenvolvimento Científico e Tecnológico, Brazil (CNPq) and by Coordenação de Aperfeiçoamento de Pessoal de Nível Superior (CAPES). GAR holds a Canada Research Chair in Genetics of the Nervous System and the Wilder Penfield Chair in Neurosciences.

