## Supplementary Material for "Genome-wide association study identifies genetic factors that modify age at onset in Machado-Joseph disease"

**Supplementary Table 1.** Linear relationship between AO and CAG_exp_, CAG_nor_, geographical origin, sex and pairwise interaction of the given factors. A total of 62.7 % of the variability in the AO is explained by given factors.

| Model description | Multiple R^2^ | Adjusted R^2^ | P-value | ΔR^2^ |
| --- | --- | --- | --- | --- |
| AO ~ CAG_exp_ | 0.6200 | 0.6195 | <2.2 × 10^−16^ |  |
| AO ~ CAG_exp_ + origin | 0.6241 | 0.6216 | <2.2 × 10^−16^ | 0.0021 |
| AO ~ CAG_exp_ + origin + sex | 0.6265 | 0.6235 | <2.2 × 10^−16^ | 0.0019 |
| AO ~ CAG_exp_ + origin + sex + CAG_nor_ | 0.6282 | 0.6247 | <2.2 × 10^−16^ | 0.0012 |
| AO ~ CAG_exp_ + origin + sex + CAG_nor_ + CAG_exp_:CAG_nor_ | 0.6301 | 0.6261 | <2.2 × 10^−16^ | 0.0014 |
| AO ~ CAG_exp_ + origin + sex + CAG_nor_ + CAG_exp_:CAG_nor_ + CAG_exp_:origin | 0.6328 | 0.6267 | <2.2 × 10^−16^ | 0.0006 |
| AO ~ CAG_exp_ + origin + sex + CAG_nor_ + CAG_exp_:CAG_nor_ + CAG_exp_:origin + CAG_nor_:origin | 0.6352 | 0.6271 | <2.2 × 10^−16^ | 0.0004 |

**Supplementary Table 2.** Subjects and cohort demographics. M:F - male-female ratio

| Geographical origin | # of patients | Mean (SD) AO | M:F |
| --- | --- | --- | --- |
| Portugal | 330 | 40.0 (±12.4) | 1.0 |
| Brazil | 311 | 34.9 (±11.7) | 1.1 |
| North America | 55 | 37.8 (±12.2) | 0.7 |
| Germany | 51 | 37.6 (±9.2) | 1.2 |
| NA | 34 | 37.1 (±11.1) | 1.4 |
| Australia | 5 | 52.8 (±10.1) | 0.3 |

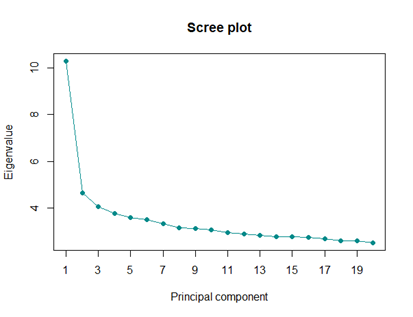

**Supplementary Figure 1.** Scree plot showing the eigenvalues of the first 20 principal components (PCs). This plot indicates that the first three PCs explain the majority of the variability in data.

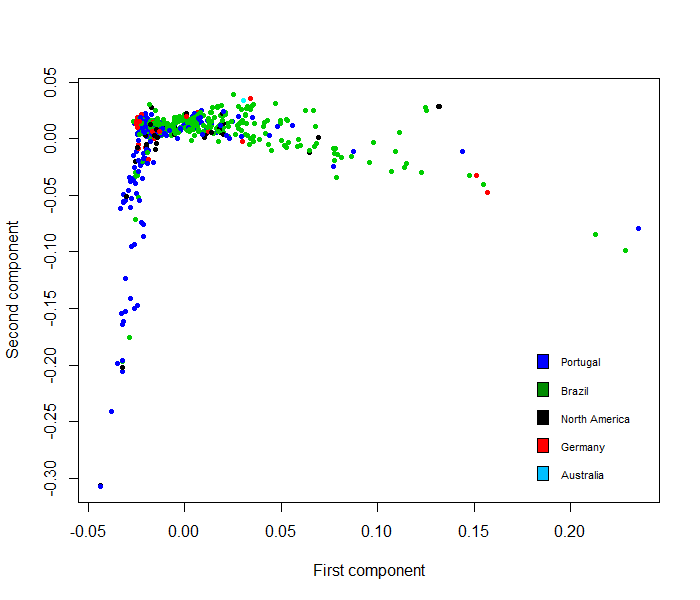

**Supplementary Figure 2.** Principal component analysis of 700 MJD patients from five different populations used for GWAS.

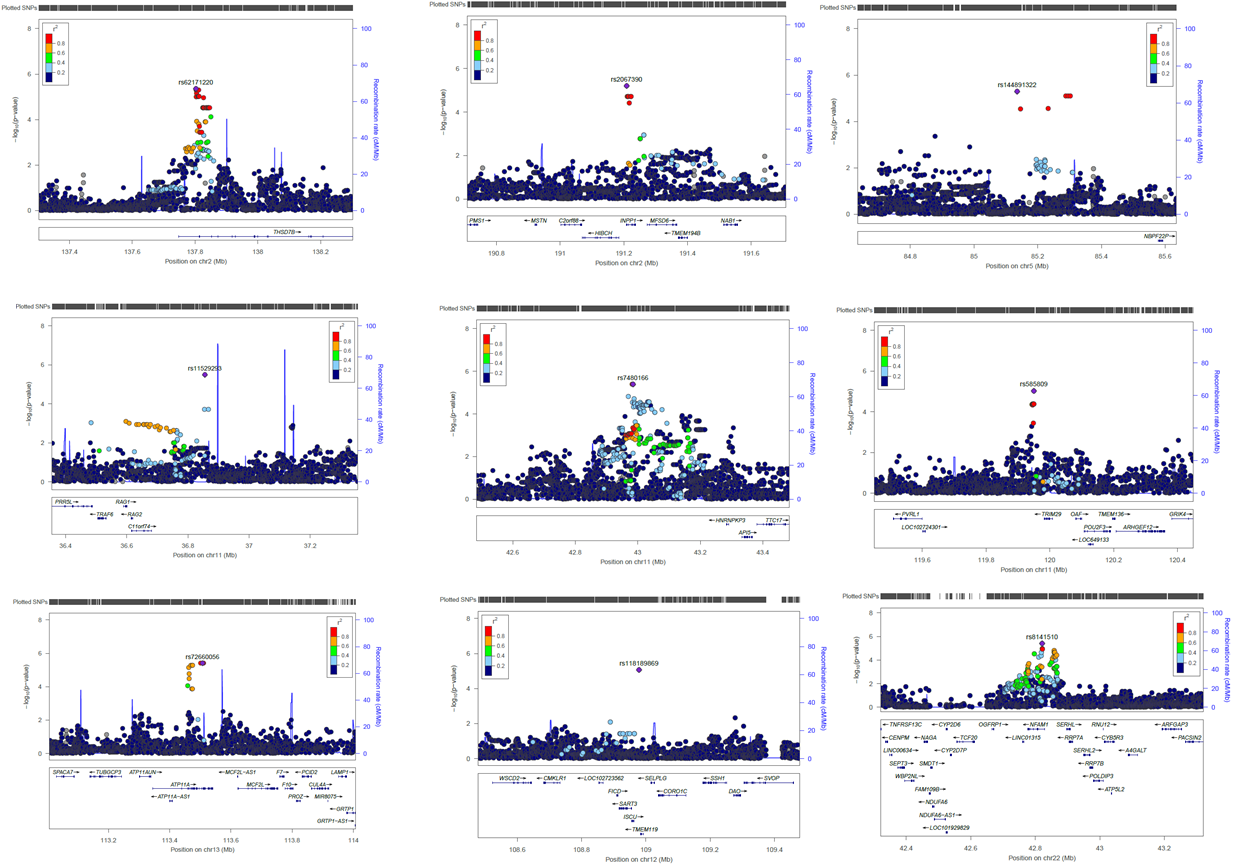

**Supplementary Figure 3.** Regional LocusZoom plots for the nine modifier loci that modify AO of MJD. Purple line indicates the genetic recombination rate (cM/Mb). SNPs in linkage disequilibrium with identified are shown in color gradient indicating r2 levels (hg19, 1KGP, Nov 2014, EUR).
